# An integrative molecular systems approach unravels mechanisms underlying biphasic nitrate uptake by plant nitrate transporter NRT1.1

**DOI:** 10.1101/2025.01.28.635294

**Authors:** Seemadri Subhadarshini, Sarthak Sahoo, Mohit Kumar Jolly, Mubasher Rashid

## Abstract

Elucidating the mechanisms of transport kinetics in plants is crucial to develop crops that can use nutrients efficiently. The plant nitrate transporter NRT1.1 rapidly switches between high- and low-affinity transport modes to maintain an optimal uptake amidst fluctuations in nitrate levels. This functional switch is regulated by NRT1.1 phosphorylation, but the precise mechanisms remain poorly understood. Here, using an integrated molecular and systems-level modeling, we identify mechanisms underlying biphasic behaviour of NRT1.1. Phosphorylation of NRT1.1 and its binding to nitrate impacts its overall flexibility and synergistically modulates its global conformation, impacting the nitrate transport rate. Integrating these observations with a regulatory network involving kinases CIPK8/CIPK23 and calcium binding proteins CBL1/9, reveals that in high nitrate conditions, CIPK8-mediated sequestration of CBL1 disrupts the CIPK23-CBL complex required for NRT1.1 phosphorylation, switching NRT1.1 to a low-affinity mode. Together, our findings untangle the molecular complexity enabling NRT1.1 phosphorylation switch with broader implications in nitrate sensing and molecular-level adaption to fluctuating external nutrient levels.

## Introduction

Plants require nitrogen for various physiological and biochemical processes essential for their growth and development. Nitrate, as a primary source of inorganic nitrogen, plays a crucial role not only as a nutrient but also as a signaling molecule [1]. It regulates nutrient transport, assimilation, and adaptive growth responses in plants [2]. Nitrate concentrations in soil can fluctuate widely, typically ranging from low micromolar to high millimolar levels, depending on factors such as microbial activity, soil composition, and agronomic practices [3, 4]. To cope with these variations, plants have developed nitrate transport systems with distinct kinetics [5, 6], ensuring efficient uptake even under low soil nitrate conditions and fine-tuning the uptake to prevent toxicity at high soil nitrate levels. Two primary families of nitrate transporters involved in this process are the High-Affinity Transport System (HATS) and the Low-Affinity Transport System (LATS). HATS includes members of the NRT2/NNP family, which facilitate nitrate uptake with Michaelis constant (Km) values in the micromolar range, making them highly effective at low nitrate concentrations. Conversely, LATS comprises the NRT1/PTR family, which transports nitrate at millimolar concentrations, providing a higher capacity for uptake when nitrate is abundant [7, 8].

Among the NRT1/PTR family members, NRT1.1, also known as CHL1 or NPF6.3, is a notable exception, distinguished by its unique ability to exhibit dual affinity for nitrate uptake [9, 10]. Identified first in *Arabidopsis thaliana* [11], this transporter can switch between high-affinity and low-affinity modes depending on the external nitrate concentration and phosphorylation status [12]. Under low nitrate conditions (< 1 mM), NRT1.1 is phosphorylated at the threonine residue 101, which enhances its affinity for nitrate. At higher nitrate concentrations (> 1 mM), the transporter remains unphosphorylated, thereby facilitating uptake in its low-affinity mode [3, 13]. Using heterologous Xenopus oocyte expression system, it has previously been demonstrated that NRT1.1 displays biphasic kinetics, characterized by two distinct Km values: ∼50 µM for the high-affinity phase of nitrate uptake and ∼4 mM for the low-affinity phase [14, 15]. This dual-affinity feature enables plants to adapt efficiently to varying nitrate levels in the soil, optimizing their nutrient acquisition and utilization [16]. Furthermore, the activity of these transport systems can be fine-tuned through transcriptional regulation and post-translational modifications, in response to changes in the nitrogen availability and the metabolic demands of the plants, enabling them to dynamically manage their nutrient uptake and maintain homeostasis. Understanding how plants perceive nitrate and transduce this signal into adaptive responses that optimize nitrate uptake via a complex interplay of transcriptional and post-translational mechanisms, particularly in the context of NRT1.1, is crucial for advancing our knowledge of plant nutrient acquisition and uptake [16, 17]. What remains largely undetermined is how the molecular structure and dynamics of NRT1.1 are altered by phosphorylation and dephosphorylation to switch between its high- and low-affinity states. Additionally, the signal transduction pathways leading to transcriptional-level changes that underpin this biphasic response are yet to be fully elucidated.

Here, we utilize the recently resolved structure of NRT1.1 [3, 13] to provide the structural basis to the role of phosphorylation in nitrate uptake. Our analysis reveals that phosphorylation at Threonine 101 significantly enhances the structural flexibility of NRT1.1, enabling dynamic shifts between conformational states, including transitions from outward/inward closed/open to outward/inward open/closed states. This conformational plasticity is critical for facilitating efficient nitrate uptake, particularly under conditions of low nitrate availability in the soil. Phosphorylation-induced flexibility broadens the conformational ensemble of NRT1.1, allowing the transporter to adopt the configurations that are optimized for nitrate binding and transport. To investigate how phosphorylation and conformational dynamics influence the rate of nitrate transport, we developed a mathematical model of ion transport based on a well-established framework. This model considers a series of reversible transitions between outward-open, occluded, and inward-open states to transport ions across the membrane. In our simulations phosphorylation reduces the conformational transition rate of NRT1.1. This reduction in transition rate likely prioritizes substrate retention, thereby enhancing transport efficiency under low-nitrate conditions. The increased flexibility of the phosphorylated form enables the transporter to achieve a trade-off between transport rate and substrate affinity, aligning its functionality with the nitrate availability in the environment. These findings underscore the critical role of structural flexibility in fine-tuning the biphasic transport behaviour of NRT1.1. Finally, to unravel the molecular determinants of phosphorylation and its connection with the biphasic regulation of NRT1.1, we developed a mathematical model of a NRT1.1-mediated nitrate-signaling network that integrates transcriptional as well as posttranslational regulators of NRT1.1. We show that while preferential binding of CBL1/9 to CIPK8 over CIPK23 is a key determinant of the phosphorylation status of NRT1.1, the biphasic expression is however independent of the phosphorylation. This approach combines detailed structural analysis with systems-level modeling to provide a holistic understanding of the regulation of this membrane transporter in nitrate uptake. Our results deciphering these intricate regulatory mechanisms have implications in developing targeted strategies to improve nutrient use efficiency of crops, ultimately supporting sustainable agricultural practices and food security.

## Results

### Threonine 101 phosphorylation elicits substantial structural and flexibility alterations in the apo form of NRT1.1 extending beyond the local site

NRT1.1 is a transmembrane protein that spans the plasma membrane of plant root cells. The gene encoding NRT1.1 in *Arabidopsis Thaliana* produces a 590-amino-acid protein, which is a member of the major facilitator superfamily (MFS) of secondary active transporters. These transporters rely on protons to transport nitrate and switch between outward and inward facing conformation states to transport ions across the membrane [3]. In its functional form, NRT1.1 crystallizes as a homodimer, with two monomers positioned side-by-side. The structure of NRT1.1 has been determined in the inward-open conformation (**Figure 1A**). Each monomer is composed of 12 transmembrane (TM) helices, organized into two distinct bundles of six helices each. These bundles form a central channel that spans the membrane, with a pseudo two-fold axis aligning the amino-terminal (TM 1-6) and carboxy-terminal (TM 7-12) domains (**Figure 1A, ii**). The two substrate-transporting tunnels, one within each monomer, are not parallel to the central axis but instead slant at an angle in opposite directions, diverging slightly from each other (**Figure 1 A, iii**). Each tunnel contains a nitrate-binding site, essential for specific recognition and binding of nitrate ions and a phosphorylation site, critical for regulation of the transport activity. Previous biochemical data showed that NRT1.1 is a dual-affinity transporter and in low nitrate conditions, phosphorylation at Threonine 101 is responsible for the switch from low- to high-affinity uptake mode to adapt to the availability of resources in the root environment [14, 18], but how phosphorylation regulates transport is largely undetermined. While some recent studies hint that phosphorylation might increase transport because of structural modifications [13, 19], a clear understanding of the impact of phosphorylation on the flexibility of NRT1.1 and its connection with transport rates is still missing.

**Figure 1:**
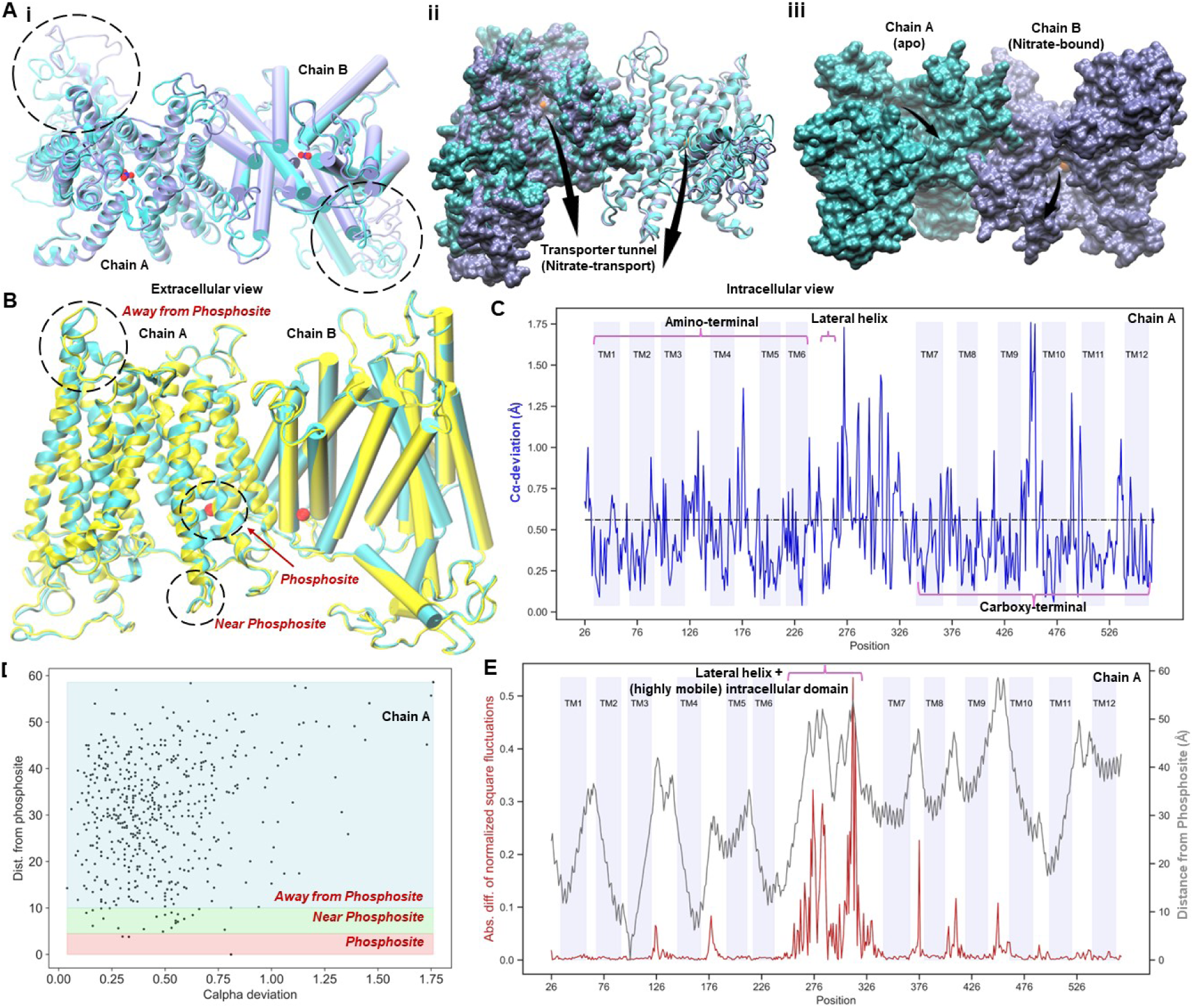
Comparative Structural Dynamics Analysis of NRT1.1 wildtype apo and T101 phosphorylated forms. **(A) (i)** Extracellular view of structural superimposition and alignment of the nitrate transporter NRT1.1 in its wildtype apo form (PDB ID: 5a2n, coloured light blue) and the nitrate-bound NRT1.1 (PDB ID: 5a2o, coloured violet). The nitrate molecule is visualized as Van der Waals spheres, with nitrogen atoms depicted in blue and oxygen atoms in red. Chain A of NRT1.1 is illustrated using “new cartoon” representation, while Chain B is depicted as “Cartoon.” The black circles indicate regions exhibiting substantial structural disparities. **(ii)** Superimposed apo and nitrate-bound forms of NRT1.1, with chain A depicted as a surface representation and chain B represented as a “new cartoon” representation. **(iii)** Surface representation of NRT1.1 with chain A of the apo form and chain B of the nitrate-bound form. The nitrate molecule is coloured orange, and arrows indicate the transporter tunnel. Both (ii) and (iii) provide an intracellular perspective. **(B)** Structural superimposition of the apo form and the threonine 101 phosphorylated form of NRT1.1, with the latter coloured yellow. Black circles highlight regions with structural disparities, including those at the phosphosite, regions near the phosphosite, and regions away from the phosphosite. The red sphere represents the phosphorylated residue. **(C)** Cα deviations between the threonine 101 phosphorylated and apo forms of NRT1.1 for all aligned residues (chain A), indicated by a blue line. The overall root mean square deviation (RMSD) between these two forms is denoted by a black line. Regions corresponding to transmembrane helices (TM1 to TM12) are coloured lavender. **(D)** Scatterplot between Cα deviations and the distance of corresponding residues from the phosphosite. Regions labelled as “Phosphosite,” “Near Phosphosite,” and “Away from Phosphosite” are demarcated by red, green, and blue shades, respectively, in the plot. **(E)** Absolute difference between normalized square fluctuations of the threonine 101 phosphorylated and apo forms of NRT1.1 for all aligned residues (chain A), depicted by a maroon line. The grey line represents the distance of each residue from the phosphorylation site.

To evaluate the conformational changes resulting from phosphorylation at the Threonine 101 residue, we compared the crystal structure of the apo form of NRT1.1 (RCSB PDB ID: 5A2N) with an in-silico modelled version of its phosphorylated counterpart (5a2nP). Structural alignments between the phosphorylated and unphosphorylated apo forms of NRT1.1 revealed alterations, not just at the phosphorylation site, but also in regions both proximal and distal to it (**Figure 1B**). To quantitatively assess the conformational differences between the two forms, we calculated the inter-residue Cα-deviation for all structurally aligned residues for both Chain A and Chain B of the NRT1.1 homodimer. We also computed the overall RMSD between the structures, with the resulting deviations represented by the blue line and the RMSD by the black line (**Figure 1C, Supplementary Figure 1A, i**). To pinpoint the regions contributing most significantly to these deviations, we created a 2D scatter plot, where the x-axis represents the Cα-deviation of each residue, and the y-axis indicates the distance of each residue from the phosphorylation site at Threonine 101. Our analysis revealed that residues with the highest Cα-deviations are predominantly located farther from the phosphorylation site (**Figure 1D, Supplementary Figure 1A, ii**). This observation suggests that phosphorylation at Threonine 101 induces allosteric conformational changes that are propagated to distant regions of the protein, potentially influencing its overall structure and function.

Protein flexibility plays a crucial role in facilitating various biological functions, as it allows proteins to adopt multiple conformations necessary for processes such as ligand binding, enzyme catalysis, and signal transduction. Understanding long-range fluctuations within a protein is particularly important, as these movements often correspond to functionally relevant motions that are essential for the protein’s activity. To study such long-range fluctuations, Normal Mode Analysis (NMA) is frequently employed [20–23]. NMA is a powerful computational technique used to explore the intrinsic motions of a protein, capturing its collective movements. To investigate the dynamic shifts resulting from phosphorylation, we performed an anisotropic network model-based normal mode analysis (ANM-NMA) on the phosphorylated and unphosphorylated apo forms. Specifically, we computed the absolute difference in inter-residue normalized square fluctuations between the two forms. These square fluctuation differences were then plotted as a function of topologically equivalent positions derived from sequence alignment (shown by the red line), as well as the distance of residues from the phosphorylation site (depicted by grey line). Our analysis revealed significant fluctuations and increased mobility, particularly within the lateral helix and the intracellular domain regions that connect the amino-terminal and carboxy-terminal domains (**Figure 1E, Supplementary Figure 1A, iii**). This observation suggests that phosphorylation at Threonine 101 not only induces localized conformational changes but also influences the dynamic behaviour of distant regions, potentially impacting the protein’s overall function.

### Post-phosphorylation and binding to nitrate, NRT1.1 undergoes extensive structural modulation, accompanied by dynamic alterations and perturbations in low-frequency global modes

To explore the cumulative effect of nitrate binding and Threonine 101 phosphorylation on the NRT1.1 homodimer, we first modelled the phosphorylation at the Threonine 101 residue position on the NRT1.1 crystal structure bound to nitrate (RCSB PDB ID: 5A2O) in silico (see Methods). We then structurally aligned this phosphorylated form (5a2oP) with the wild type unphosphorylated apo form (RCSB PDB ID: 5A2N). Our analysis revealed deviations not only at the phosphorylation site but also in regions both near and distant from it (**Figure 2A, i**). The NRT1.1 transporter spans the cellular membrane, with regions that extend outward towards the extracellular space and inward into the cytosol. The regions of the protein that lie towards the cytosolic side exhibit significant structural disparities between the phosphorylated nitrate-bound and unphosphorylated forms (**Figure 2A, ii**). The three crystal structures of the nitrate transporter, with RCSB PDB IDs 5A2N and 5A2O [13], and 4OH3 [3], were used as a basis for modelling their phosphorylated counterparts. We then estimated the Cα RMSD as a metric to understand the conformational disparity induced by phosphorylation. Additionally, we computed the Cα RMSD for phosphorylated nitrate-bound form in comparison to the unphosphorylated apo form to assess the conformational changes resulting from the combined effects of phosphorylation and nitrate binding. Our analysis revealed that the Cα RMSD increases by approximately 6-fold, when comparing phosphorylated nitrate-bound form to the unphosphorylated apo form, as opposed to comparing just the phosphorylated forms to the unphosphorylated apo form (**Figure 2B, Supplementary Figure 2, Supplementary Figure 3**). This observation indicates that the combined effect of phosphorylation and nitrate binding induces more pronounced conformational changes in the transporter.

**Figure 2:**
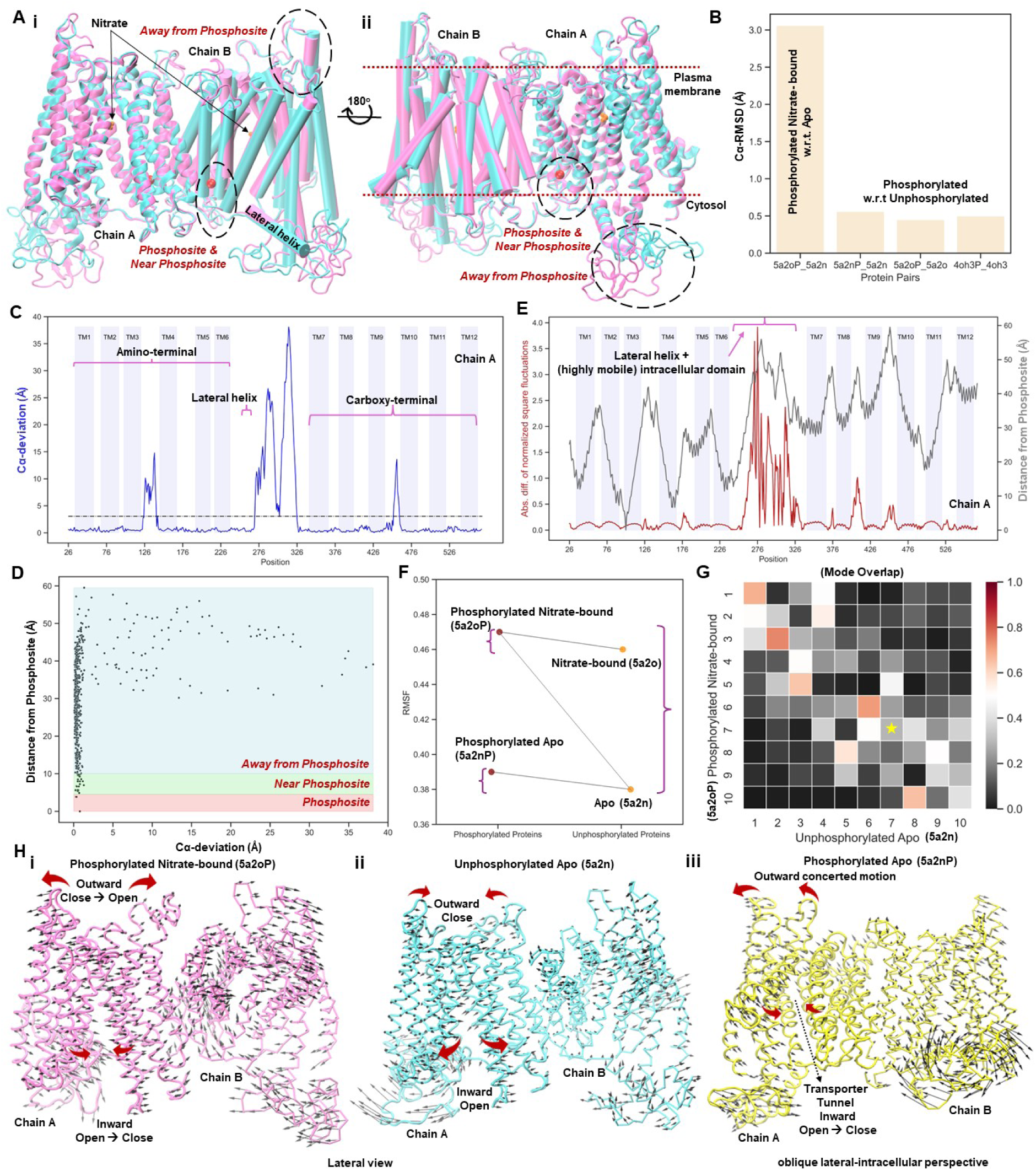
Differential conformational changes highlighting localized alterations post phosphorylation and nitrate binding. **(A) (i)** Structural superimposition and alignment of the nitrate-bound T101 phosphorylated form of NRT1.1, highlighted in pink, alongside the wildtype apo form, depicted in light blue. Black circles delineate regions exhibiting structural disparities, encompassing those at the phosphosite, in proximity to the phosphosite, and distant from the phosphosite. The red sphere denotes the phosphorylated residue, while the orange sphere represents the nitrate molecule. **(ii)** In a 180-degree flipped lateral view of (i), regions of the membrane transporter facing towards the cytosol and those oriented towards the extracellular side of the plasma membrane are indicated by a delineating red line. **(B)** Barplot showing the Cα-RMSD between protein pairs: Phosphorylated nitrate-bound NRT1.1 (5a2oP) with apo form (5a2n), Phosphorylated apo (5a2nP) with apo form (5a2n), Phosphorylated nitrate-bound (5a2oP) with nitrate-bound (5a2o), and Phosphorylated nitrate-bound (4oh3P) with nitrate-bound (4oh3). **(C)** Cα deviations between the nitrate-bound, threonine 101 phosphorylated, and apo forms of NRT1.1 for all aligned residues (chain A), depicted by a blue line. The overall root mean square deviation (RMSD) between these two forms is denoted by a black line. Regions corresponding to transmembrane helices (TM1 to TM12) are coloured lavender. **(D)** Scatterplot illustrating the relationship between Cα deviations and the distance of corresponding residues from the phosphosite. Regions labelled as “Phosphosite,” “Near Phosphosite,” and “Away from Phosphosite” are demarcated by red, green, and blue shades, respectively, in the plot. **(E)** Absolute difference between normalized square fluctuations of the nitrate-bound threonine 101 phosphorylated and apo forms of NRT1.1 for all aligned residues (chain A), represented by a maroon line. The grey line represents the distance of each residue from the phosphorylation site. **(F)** Strip plot displaying RMSF values for corresponding phosphorylated and unphosphorylated proteins. **(G)** Overlap of the first ten non-zero normal modes of nitrate-bound phosphorylated and unphosphorylated forms of NRT1.1. **(H) (i)** Backbone fluctuations depicted by eigen vectors of the nitrate-bound phosphorylated form of NRT1.1. **(ii)** Backbone fluctuations depicted by eigen vectors of the unphosphorylated apo form of NRT1.1. **(iii)** Backbone fluctuations depicted by eigen vectors of the phosphorylated apo form of NRT1.1.

As before, we computed the Cα-deviation for all structurally aligned residues in the phosphorylated nitrate-bound form relative to the unphosphorylated apo form. The deviations are represented by the blue line, and the overall RMSD is shown by the black line (**Figure 2C, Supplementary Figure 1B, i**). The 2D scatter plot, with the x-axis representing the Cα-deviation of each residue and the y-axis indicating the distance of each residue from the phosphorylation site at Threonine 101, revealed that residues with the highest Cα-deviations are predominantly located farther from the phosphorylation site (**Figure 2D, Supplementary Figure 1B, ii**). Notably, these deviations are more pronounced than those observed in the solely phosphorylated forms, highlighting the added impact of nitrate binding in conjunction with phosphorylation. Next, to assess changes in the dynamics and flexibility of the protein backbone, we computed the absolute difference in inter-residue normalized square fluctuations between the phosphorylated nitrate-bound form and the unphosphorylated form, following the same approach as before. Our analysis revealed increased mobility, particularly within the lateral helix and the intracellular domain regions that connect the amino-terminal and carboxy-terminal domains. However, this increased mobility was observed to be several folds higher than in the case of phosphorylation alone (**Figure 2E, Supplementary Figure 2B, iii**), underscoring the significant impact of the combined effects of phosphorylation and nitrate binding on the structural dynamics of the transporter. Upon further scrutiny, we observed that the phosphorylated apo form exhibits moderately higher root-mean-square fluctuation (RMSF) compared to its unphosphorylated counterpart, while the phosphorylated nitrate-bound form shows even higher RMSF values **(Figure 2F**).

Low-frequency global modes derived from normal mode analysis (NMA) are crucial indicators of biologically relevant dynamics in proteins, providing key insights into their functional motions, such as enzyme-substrate binding [24], and conformational changes [20, 25, 26]. Deciphering whether the combined effects of phosphorylation and nitrate binding alter the low-frequency modes of proteins and how these modifications relate to functional dynamics is crucial for understanding the nitrate transporter. Specifically, it’s important to determine whether these changes introduce new low-frequency motions or preserve the inherent dynamics of the protein. To explore this, we analysed the similarities and differences between the modes of motion accessible to the protein in its phosphorylated nitrate-bound and unphosphorylated forms. This was achieved by quantifying the overlap between the top 10 low-frequency global motions obtained from NMA, where the overlap value serves as a robust indicator of similarity in frequency, shape, and size between the modes. A smaller overlap value indicates a larger difference between the two modes of motion. The low-frequency global modes showed little overlap between the phosphorylated nitrate-bound and unphosphorylated forms, with some modes displaying shifts in mode preference and order upon phosphorylation and nitrate binding. A shift in mode preference suggests that a mode present as a low-frequency mode in one form appears as the same or a similar mode (indicated by a high overlap value) in the other form but with a modified frequency. This finding implies that while a few modes of motion are somewhat preserved between the phosphorylated nitrate-bound and unphosphorylated forms, their frequency, size, and shape undergo substantial changes, as evidenced by the reordering of normal modes—though most modes are lost, indicating extensive dynamic rearrangements occurring because of nitrate binding and phosphorylation (**Figure 2G**). To further elucidate the changes in the dynamics and flexibility of the protein backbone, we closely examined the backbone motions of the NRT1.1 transporter under three distinct scenarios: (1) phosphorylated and nitrate-bound form (5a2oP), (2) unphosphorylated apo form (5a2n), and (3) phosphorylated apo form (5a2nP). We focused our analysis on Normal Mode 7, which was selected due to its alignment with the key functional motions observed in preliminary assessments (**Supplementary Movie**). All structures analysed were initially resolved in the inward-open conformation. Our analysis revealed that in the phosphorylated and nitrate-bound form (5a2oP), the transporter undergoes an outward movement from a closed to an open state on the extracellular side and an inward movement from an open to a closed state on the intracellular side (**Figure 2H, i**). This bidirectional motion is consistent with the functional requirements for nitrate transport, facilitating the transition between inward-open and outward-open states essential for substrate translocation. In contrast, the unphosphorylated apo form (5a2n) exhibited an outward closed to inward open motion (**Figure 2H, ii**). Given that this form is already in an inward-open conformation, the observed motion results in an even more open state, enhancing the accessibility of the transport tunnel. This dynamic flexibility is crucial for the regulation of nitrate transport, as the opening and closing of the tunnel on both extracellular and intracellular sides are fundamental for substrate binding and release. For the phosphorylated apo form (5a2nP), the transporter primarily transitions from an inward-open to a closed state on the intracellular side, while the extracellular side displays a concerted motion that does not fully transition to an outward-open state (**Figure 2H, iii**). This partial movement suggests that phosphorylation alone may prime the transporter for functional dynamics, potentially stabilizing certain conformational states and modulating the overall flexibility of the protein. These observations indicate that phosphorylation, especially when combined with nitrate binding, significantly alters the dynamic behaviour of NRT1.1. The phosphorylated nitrate-bound form demonstrates enhanced and more coordinated conformational changes compared to the phosphorylated apo and unphosphorylated forms, underscoring the importance of post-translational modifications in regulating transporter function. The phosphorylation-induced priming could enhance the binding affinity by stabilizing specific conformations that favour nitrate recognition but reduce the rate of transport due to incomplete structural transitions required for efficient transport. In contrast, the unphosphorylated state, which shows a more open inward-facing conformation, may support a higher rate of transport but with lower affinity, as the protein is more dynamically open, allowing for faster substrate exchange but weaker binding. This dynamic interplay between phosphorylation states likely regulates the transporter’s functional efficiency, balancing nitrate uptake and responsiveness to cellular signals.

### Phosphorylation modulates nitrate transport kinetics and biphasic behaviour of NRT1.1 by altering the inward-outward conformation transition rate

Following a detailed analysis of the structural dynamics post-phosphorylation, we aimed to investigate the differences in nitrate transport kinetics between the phosphorylated and unphosphorylated forms of NRT1.1. Transport processes across the membrane have been modelled previously using ordinary differential equations [27, 28]. We used previously developed mathematical models to study the changes in transport kinetics due to phosphorylated and unphosphorylated forms of NRT1.1 (See Methods)[28]. Briefly, the phosphorylated/unphosphorylated forms of the proteins were assumed to have 4 distinct conformational states – outward open apo form, outward open nitrate bound, inward open nitrate bound and inward open apo states (**Figure 3A**). All transitions between these 4 states are allowed albeit at different rates (**Figure 3A**). The rates involved in transitions between the outward and inward conformations were different for the phosphorylated and the unphosphorylated cases (see Methods section for modelling details and information about the parameters). Rate affinity trade-off in such systems has been previously investigated especially in the context of glucose transporters [28]. We investigated if such rate affinity trade-off is also seen in the context of NRT1.1 as a function of the phosphorylation status.

**Figure 3:**
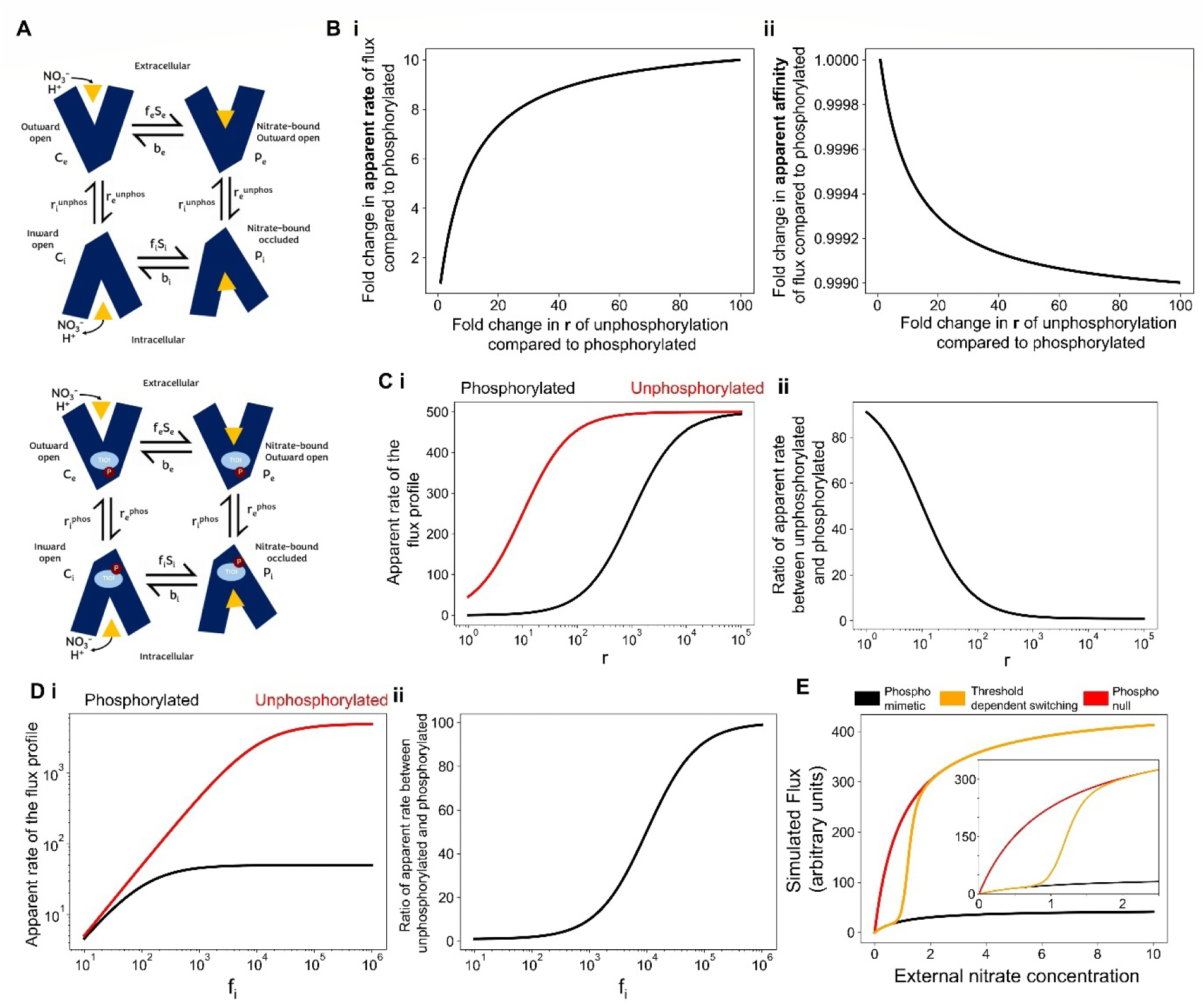
Changes in flux and associated apparent rate of transport as a function of the substrate concentration for the phosphorylated and unphosphorylated states. **A)** 4 state diagrams for the phosphorylated and the unphosphorylated forms of NRT1.1 showing the various transitions possible and the associated transition rates between the four conformations states. **B)** Line plot showing the functional relationship between the fold change of the i) apparent rate and ii) apparent affinity of transport of nitrate between the phosphorylated and unphosphorylated forms of NRT1.1. **C)** i) Line plot showing the functional relationship between the apparent rate of the flux profile with absolute value of the rate parameter r for the phosphorylated (red) and unphosphorylated (black) forms of NRT1.1. ii) Line plot showing the fold change differences between the phosphorylated and the unphosphorylated forms of NRT1.1 over multiple orders of magnitude of the parameter r. **D)** i) Line plot showing the functional relationship between the apparent rate of the flux profile with absolute value of the parameter f_i_ for the phosphorylated (red) and unphosphorylated (black) forms of NRT1.1. ii) Line plot showing the fold change differences between the phosphorylated and the unphosphorylated forms of NRT1.1 over multiple orders of magnitude of the parameter f_i_. **E)** Simulated flux of nitrate transport over different external nitrate levels in three variants of NRT1.1 – phospho-null NRT1.1 (red), phospho-mimetic (black) and wild type biphasic NRT1.1 (orange). Zoomed view of the switching in flux profiles has been showing in the inset.

Since the dissociation constant K_D_ for nitrate binding is approximately the same for both the phosphorylated and unphosphorylated forms of NRT1.1 [13], we reasoned that phosphorylation does not affect the equilibrium affinity of nitrate binding. Thus, differences in the Michaelis-Menten constant (Km) between the high- and low-affinity states must arise from other factors, likely linked to the conformational dynamics of the transporter. These dynamics could, in turn, influence the overall transition rates of the transporter between inward- and outward-facing conformations. Based on this rationale, we assumed that the key free parameter affected by phosphorylation is the rate of transition between these conformations. To distinguish between the unphosphorylated and phosphorylated forms of the transporter, we introduced distinct variables: *r^unphos^* and *r^phos^* representing the transition rates for the unphosphorylated and phosphorylated forms, respectively. Upon simulating the system of differential equations for varying relative transition rate *r* (r^unphos^/r^phos^) while keeping all other parameters fixed, we observe that an increase in this ratio - indicating a higher transition rate for the unphosphorylated state relative to the phosphorylated state - led to an increase in the apparent transport rate but a corresponding decrease in affinity (**Figure 3B, i-ii**). This result demonstrates the existence of rate-affinity trade-off linked to the phosphorylation state of NRT1.1. However, the magnitude of decrease in the affinity is minimal and might not have functional consequence (**Figure 3B, ii**). On the other hand, the increase in the apparent rate of transport significantly depends on the parameter *r* and saturates at very high levels of *r* in the unphosphorylated state compared to the phosphorylated form (**Figure 3B, i**).

We next visualized the estimated apparent nitrate transport rates as a function of the absolute values of *r^phos^ and r^unphos^ across* several orders of magnitude, while keeping the fold change between *r^phos^ and r^unphos^* constant. The analysis revealed that at very low magnitudes of *r^unphos^*, the apparent rate of nitrate transport is minimal, increases with higher values of *r^unphos^*, and eventually saturates (**Figure 3C, i-ii**). A similar trend is observed for the phosphorylated form, although the apparent transport rate for the phosphorylated state reaches 50% of its maximal value at a much lower *r^phos^* compared to the unphosphorylated state, which requires a higher *r^unphos^* to achieve the same threshold (**Figure 3C, i**). This trend indicates that the fold change in the apparent transport rates between the unphosphorylated and phosphorylated forms - i.e., the relative transport rate - is less pronounced at higher absolute values of the transition rate parameter *r*. Conversely, the difference in transport rates between the two forms is more pronounced at lower *r* (**Figure 3C, ii**). This might have implications for the absolute value of the parameter *r* at which NRT1.1 operates under physiological conditions and suggests that its functional behaviour may be tuned by the dynamic range of the rate parameter. One other parameter that can influence the rate of nitrate transport through NRT1.1 is *f_i_* which represents the association rate of the intracellular substrate to the transporter. An increase in *f_i_* leads to a corresponding increase in the apparent transport rate for nitrate (**Figure 3D, i**). This trend is qualitatively mirrored by the phosphorylated form of NRT1.1, alike its unphosphorylated state (**Figure 3D, i**). Furthermore, the fold change in the apparent transport rate between the unphosphorylated and phosphorylated forms is most pronounced at higher *f_i_* values. This indicates that the differences in nitrate uptake rates between the two forms of NRT1.1 become more evident at higher *f_i_* values (**Figure 3D, ii**).

Finally, we modelled the biphasic nature of nitrate transport through NRT1.1 (orange profile) and compared it to the simulated profiles representing phospho-null (no phosphorylation) and phospho-mimetic mutation states (red and black profiles, respectively) (**Figure 3E**). Our simulations revealed that altering only the transition rate parameter *r* between the phospho-null and phospho-mimetic cases resulted in drastic changes in the simulated nitrate flux across varying external nitrate concentrations. This observation supports our initial hypothesis that changes in conformational dynamics - specifically, altered flexibility modulating *r*, plays a critical role in driving the biphasic response of NRT1.1 to external nitrate levels. Furthermore, we could simulate the biphasic nature of the NRT1.1 switching between the two modes by changing the parameter *r* dynamically around the known threshold external nitrate concentration of 1mM [3, 13, 14] (**Figure 3E**). Our simulation results show that transport of nitrate through NRT1.1 is primarily driven by changes in the conformational transition rate parameter *r*. Importantly, gradual modulation of this parameter was sufficient to recapitulate the characteristic biphasic flux observed previously in literature [14], validating the mechanistic link between the transporter’s conformational dynamics and its nitrate transport efficiency.

### Systems-level integration of transcriptional, protein-protein interaction and post-translational modifications explain nitrate-dependent shifts in NRT 1.1 expression and phosphorylation levels

NRT1.1 functions not only as a nitrate transporter but also as a sensor, regulating its own expression and that of other nitrate-responsive genes [29, 30]. This dual functionality allows NRT1.1 to adapt to varying nitrate concentrations in the environment, effectively switching between high-affinity and low-affinity transport modes based on both external nitrate concentrations and post-translational modifications. Investigating the molecular mechanisms behind nitrate uptake and signaling that drive this biphasic mode of nitrate uptake is crucial for advancing our understanding of mineral nutrition in plants and gain insights into how plants optimize nutrient acquisition in response to fluctuating environmental conditions [31]. To explore the underlying mechanisms from a systems-level perspective, we constructed a comprehensive network model that captures this behaviour by integrating transcriptional regulation, protein-protein interactions, and post-translational modifications (**Figure 4A**). The model reveals how low external nitrate concentrations favour phosphorylated NRT1.1, while higher nitrate levels shift the balance toward its unphosphorylated form, thereby approximating the biological system at hand while providing insights into its dynamic regulation.

**Figure 4:**
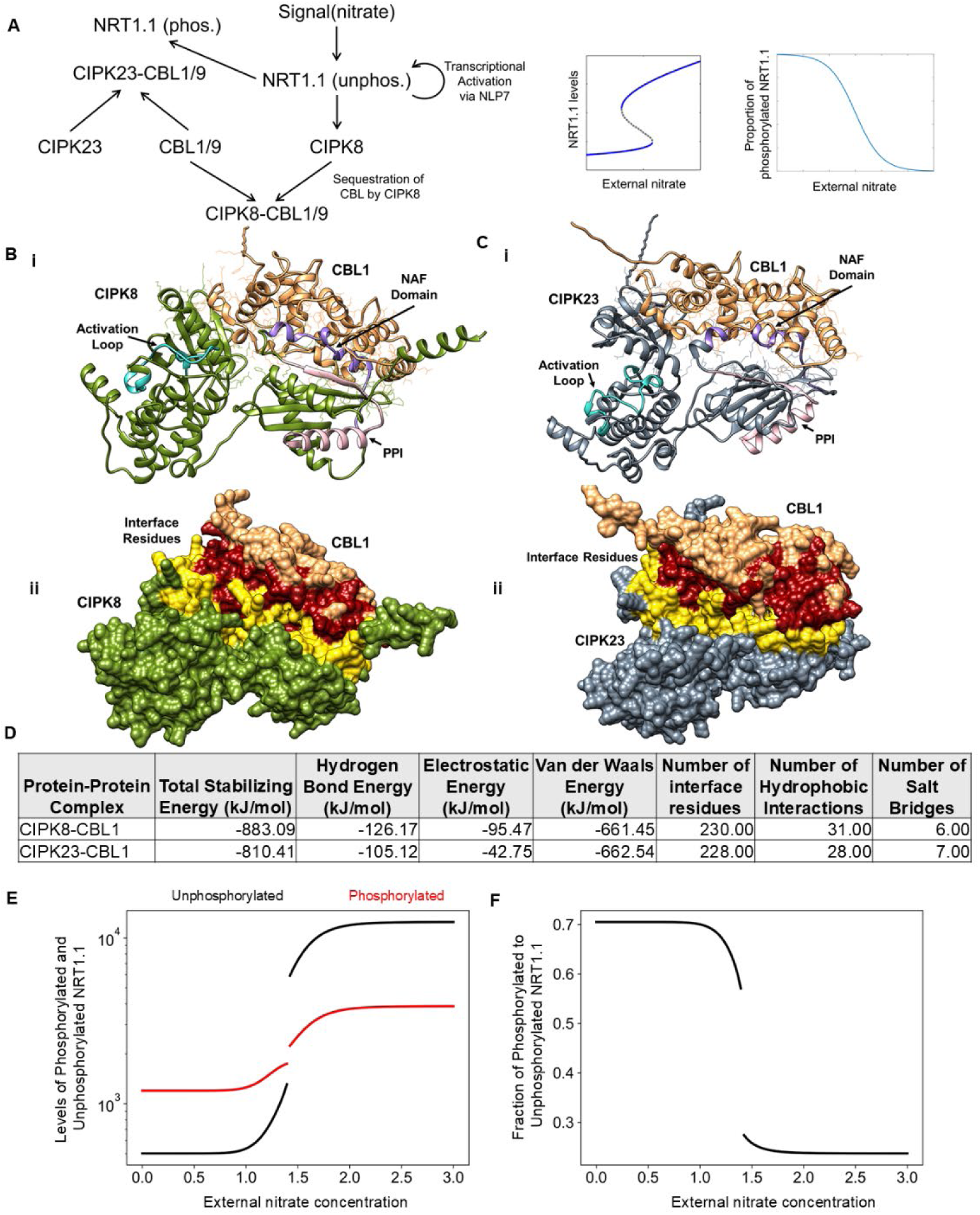
Regulatory Network Dynamics of Nitrate Response. **(A)** Network depicting nitrate-dependent shifts in NRT1.1 expression and phosphorylation, integrating transcriptional regulation, CBL1/9 sequestration by CIPK8/23, and post-translational modifications. **(B) (i)** Docked CIPK8-CBL1 Complex: CIPK8 is depicted in olive, and CBL1 is shown in light brown, with both protein backbones represented as ribbons. The side chains of interfacial residues between the CIPK8-CBL1 complex are displayed as sticks. The activation loop of the kinase is coloured light blue, the NAF domain is purple, and the PPI region is pink. **(ii)** Surface Representation of the CIPK8-CBL1 Docked Complex: The surface of the kinase CIPK8 is coloured olive, and the surface of CBL1 is light brown. Interfacial residues of CIPK8 are highlighted in yellow, while those of CBL1 are shown in red. **(C) (i)** Docked CIPK23-CBL1 Complex: CIPK23 is depicted in dark grey, and CBL1 in light brown, with both protein backbones represented as ribbons. The side chains of interfacial residues in the CIPK23-CBL1 complex are displayed as sticks. The activation loop of the kinase is coloured light blue, the NAF domain is purple, and the PPI region is pink. **(ii)** Surface Representation of the CIPK23-CBL1 Docked Complex: The surface of CIPK23 is coloured dark grey, and the surface of CBL1 is light brown. Interfacial residues of CIPK23 are highlighted in yellow, while those of CBL1 are shown in red. **(D)** Comparative table showing total stabilizing energy, hydrogen bond energy, electrostatic energy, van der Waals energy, number of interface residues, hydrophobic interactions, and salt bridges for CIPK8-CBL1 and CIPK23-CBL1 complexes. **(E)** Simulated levels of phosphorylated (black) and unphosphorylated (red) forms of NRT1.1 as a function of external nitrate concentration at steady state. **(F)** Simulated proportions of fraction of phosphorylated to unphosphorylated forms of NRT1.1 as a function of external nitrate concentration at steady states.

External nitrate acts as a signal, sensed primarily by molecular components involved in the primary nitrate response (PNR), including NRT1.1 transceptor (transporter cum sensor). The sensor function of NRT1.1 is triggered by rising nitrate levels, activating downstream signaling pathways, including transcription factors like NLP7 (NIN-like protein 7), which regulate the expression of nitrate-responsive genes, such as NRT1.1 itself. Additionally, other critical genes in nitrate signaling, like CIPK8 (CBL-Interacting Protein Kinase 8), are also upregulated as part of this response. CIPK8 and CIPK23 (Calcineurin B-Like Interacting Protein Kinases) are important components of the nitrate signaling pathway. Both proteins interact with CBL1 and CBL9 (Calcineurin B-like proteins), which are crucial for mediating calcium signaling in response to nitrate **(Figure 4A)**[29, 30, 32–34]. When nitrate levels are high, increased levels of CIPK8 can sequester CBL1/9, leading to a reduction in the availability of these proteins for interaction with CIPK23. This preferential sequestration results in a decreased formation of the CIPK23-CBL complex, which is essential for the phosphorylation of NRT1.1, leading to a decrease in NRT1.1’s phosphorylation and a shift toward its low-affinity state. Conversely, under low nitrate conditions, the CIPK23-CBL complex formation is favoured, resulting in enhanced phosphorylation of NRT1.1 and maintaining its high-affinity state. This dynamic regulation of NRT1.1 phosphorylation helps plants finely tune nitrate uptake in response to environmental nitrate levels. The interaction between CIPK (CBL-Interacting Protein Kinases) and CBL (Calcineurin B-like proteins) is crucial in modulating the phosphorylation status of NRT1.1, thereby influencing the plant’s nitrate response. CIPKs bind to CBLs through their NAF domain, a conserved region that facilitates the formation of CIPK-CBL complexes. Once bound, the CIPK-CBL complex can interact with target proteins such as NRT1.1, particularly at the PPI (Protein-Protein Interaction) region, to regulate its phosphorylation. Our analysis revealed that CBL1 binds differentially to CIPK8 and CIPK23. Upon docking CBL1 with these kinases, we observed that the CIPK8-CBL1 complex (**Figure 4B, i-ii**) is energetically more favorable than the CIPK23-CBL1 complex (**Figure 4C, i-ii**). The total stabilization energy for the CIPK8-CBL1 complex was calculated to be −883.09 KJ/mol, compared to −810.41 KJ/mol for the CIPK23-CBL1 complex (**Figure 4D**). This suggests that the CIPK8-CBL1 interaction is more stable, leading to preferential sequestration of CBL1 by CIPK8 in high nitrate conditions. As a result, fewer CBL1 molecules are available for forming the CIPK23-CBL complex, reducing the phosphorylation of NRT1.1 and keeping it in its low-affinity state. This differential binding and sequestration mechanism may play a key role in fine-tuning the nitrate response in plants.

We simulated the regulatory network using a set of coupled differential equations (see methods) to model how the levels of NRT1.1 could change as a function of systems level changes in signalling and consequent changes in its phosphorylation status. As the model parameters are not available in the existing literature, our model represents a possible mode of coordination of NRT1.1 phosphorylation status and its resultant flux in nitrate transport which would require future experimental validation. Upon simulation of the set of differential equations we observed that the NRT1.1 levels in higher nitrate concentration went up significantly (**Figure 4E**). This is consistent with previous literature showing that significant upregulation in transcription of the NRT1.1 receptor expression is seen at higher external nitrate concentrations. However, we see that although the absolute levels of both the phosphorylated and the unphosphorylated forms of NRT1.1 increase (**Figure 4E**) the proportion of the phosphorylated form of NRT1.1 drops significantly at higher levels of external nitrate concentration (**Figure 4F**). Our results show that along with significant upregulation of NRT1.1 the subsequent activation of CIPK8 can act to decrease the phosphorylation level of the NRT1.1 thus transferring most of the NRT1.1 to an unphosphorylated form which leads to overall increase in nitrate transport both due to increased individual molecules’ transport capacity as well as due to large upregulation of the number of molecules with a systems levels coordination controlling the upregulation of the overall capacity of cells to efficiently transport nitrate inside the cells.

## Discussion

The investigation into NRT1.1’s functionality as a dual-affinity transporter began with the observation that it operates with two distinct Km (Michaelis Menten constant) values [15, 35], allowing it to switch between high-affinity and low-affinity transport modes depending on the external nitrate concentration [36]. This remarkable ability enables plants to optimize nutrient uptake in varying environmental conditions. A critical finding in this context was the role of phosphorylation at the Threonine 101 residue, which was shown to facilitate this switching mechanism [37]. However, the underlying rationale for this transition between affinity states remained unclear, particularly how phosphorylation modulates the transport kinetics and expression levels of NRT1.1 in response to external nitrate availability. Previous studies have established that the expression of NRT1.1 is intricately tied to external nitrate levels, with higher concentrations promoting increased transporter expression [38]. This regulatory response suggests a sophisticated adaptation mechanism, allowing plants to fine-tune their nutrient acquisition strategies in accordance with fluctuating nutrient environments.

Here, we elucidate the molecular mechanisms that govern these observations, particularly the role of Threonine 101 phosphorylation in mediating the transporter’s switching behaviour. To address this complex issue of biphasic behaviour of NRT 1.1, we adopted an approach integrating structural analyses and kinetic modeling. At the structural level, we investigated how phosphorylation affects the protein conformation of NRT1.1, potentially altering its transport capacity and affinity. Concurrently, we examined the transport flux kinetics pre- and post-phosphorylation to understand the rate-affinity relationships that characterize this dual functionality. Finally, we explored the systems-level dynamics of downstream signaling pathways following the sensing of external nitrate levels, aiming to reveal how these pathways influence NRT1.1 expression and phosphorylation states. Our findings offer a comprehensive understanding of how plants regulate nitrate transport in response to environmental changes, shedding light on the complex interplay between signaling, post-translational modifications, and transporter functionality. We show that phosphorylation leads to distinct conformational alterations at and around Threonine 101 residue that impacts flexibility in the transporter structure and consequent protein’s dynamic behaviour. Additionally, we observed that phosphorylation influences not only local structural changes at the phosphorylation site but also global alterations within the protein, consistent with previous observations [21]. These changes may be pivotal in facilitating the transition between high-affinity and low-affinity states, as the protein must undergo substantial conformational shifts to accommodate substrate binding and transport. To further examine the underlying mechanisms governing NRT1.1’s functionality, we evaluated the transport kinetics pre- and post-phosphorylation, and observed a rate-affinity trade-off in nitrate transport. While the dissociation constant (Kd) remains constant between phosphorylated and unphosphorylated forms, the rate of transport is primarily influenced by the transition rate parameter *r*. In the phosphorylated, high-affinity state, increased flexibility reduces the transition rate between conformational states (e.g., outward-open ↔ inward-open). This slower cycling enhances substrate retention, effectively lowering Km and prioritizing efficient nitrate transport under low external nitrate conditions. Conversely, in the unphosphorylated, low-affinity state, faster conformational transitions (higher *r*) facilitate higher throughput, favouring transport at high nitrate concentrations. As *r* increases, the apparent rate of nitrate transport rises, albeit with a minimal decrease in affinity. This finding is consistent with previous observations in glucose transporters, suggesting that similar kinetic principles apply to the nitrate transporter NRT1.1 [28].

Moreover, our simulations indicated that the apparent transport rates saturate at high values of r. This trend could help cells adapt to fluctuating environmental nitrate levels by having a threshold behaviour, and also implies a limit to the efficiency of nitrate transport. The dual functionality of NRT1.1 as both a transporter and a sensor allow it to adapt to external nitrate concentrations, switching between high-affinity and low-affinity states. Our findings indicate that low external nitrate levels favour the phosphorylated form of NRT1.1 while high nitrate concentrations lead to an increased prevalence of the unphosphorylated form, promoting a shift toward low-affinity transport. Phosphorylation broadens the conformational ensemble of the transporter, enabling it to explore a wider range of states. At low nitrate concentrations, the high-affinity state prioritizes retention and efficient transport, while at high nitrate concentrations, the low-affinity state supports faster transitions and greater transport capacity. The integration of transcriptional regulation and protein interactions further elucidates how NRT1.1 regulates its own expression. The involvement of key transcription factors like NLP7 and the intricate interactions between CIPKs and CBLs highlight the nuanced signaling pathways that dictate the phosphorylation status of NRT1.1 [16, 38]. Specifically, our data show that elevated levels of CIPK8 in high-nitrate conditions sequester CBL1/9, reducing their availability to form the CIPK23-CBL complex, which is critical for NRT1.1 phosphorylation. This preferential sequestration mechanism allows plants to finely tune their nitrate uptake strategy in response to external nitrate levels. The systems-level model constructed in this study not only captures the regulatory dynamics of NRT1.1 but also reflects the broader implications of nitrate signaling in plant physiology. The balance between NRT1.1 phosphorylation states could serve as a key regulatory mechanism, enabling plants to efficiently adapt to varying nitrate levels.

While our model presents a compelling framework for understanding NRT1.1 regulation, further experimental validation to estimate the parameters corresponding to transport kinetics captured is crucial. Future studies should focus on biochemical and structural assays to determine the precise binding affinities, phosphorylation kinetics, and conformational changes associated with Threonine 101 phosphorylation. Also, time-resolved nitrate transport assays could provide insights into how phosphorylation influences NRT1.1’s biphasic transport behaviour in real-time, offering a better understanding of the rate-affinity relationship. Overall, our study represents the first integrative attempt to understand the biphasic behaviour of NRT1.1, bridging the gap between structural changes at the molecular level and systems-level nitrate signaling. These findings can serve as crucial inputs to biotechnological interventions aiming to enhance the nitrate uptake efficiency in plants to increase crop yields and reduce the pollution due to unutilized nitrogen-fertilizers.

## Methods

### Structural Analysis

#### Structural Data Selection

The RCSB Protein Data Bank (PDB) contains three reported crystal structures for the dual-affinity nitrate transporter NRT1.1 from Arabidopsis thaliana, identified by PDB IDs 5A2N, 5A2O, and 4OH3. All three structures are homodimers and represent wild type unphosphorylated variants. PDB IDs 4OH3 and 5A2O are bound to a nitrate ligand, whereas 5A2N is an apo structure (no ligand bound). The structures with PDB IDs 5A2N and 5A2O were deposited by Parker et al. [13], while 4OH3 was deposited by Sun et al. [3].Both 5A2N and 5A2O were resolved at 3.7 Å resolution and were selected as the primary structures for phosphorylation analysis. For completeness, 4OH3 was also analysed pre- and post-phosphorylation and is included in the Supplementary Information.

#### Modelling Non-Terminal Missing Residues

All selected structures had missing residues that required modelling. Non-terminal missing residues were modelled using the Modeller software, a widely used tool for comparative modelling [39]. Modeller employs iterative optimization methods to generate accurate protein models by satisfying spatial restraints and optimizing various energy functions. Fifty models were generated for each structure, each representing different conformations of the missing residues. The loop modelling protocol in Modeller, which utilizes distance geometry optimization and molecular dynamics, was employed. The quality and reliability of the generated models were assessed using the DOPE-HR (Discrete Optimized Protein Energy) energy function. DOPE-HR is a composite scoring function that evaluates the likelihood of a model being native-like based on various physical and statistical potential terms. The models were ranked according to their DOPE-HR scores, with lower scores indicating higher quality. The model with the lowest (most negative) DOPE-HR score was selected as the best representation for further analysis.

#### In-Silico Phosphorylation

The modelled structures, now complete with no missing residues, underwent in-silico phosphorylation at the threonine 101 residue in both chain A and chain B. This process was carried out using the CHARMM 36 force field and the GROMACS molecular dynamics software package [40]. To facilitate this, the solution builder module of the CHARMM-GUI input generator was employed. The CHARMM-GUI solution builder provides an intuitive interface for defining the system, adding the desired modifications such as phosphorylation, and generating the necessary input files for subsequent simulations [41]. The tool also ensures that the modified residues, including the phosphorylated threonine, are appropriately parameterized according to the CHARMM 36 force field specifications [42].

#### Energy Minimization

Following the in-silico phosphorylation, the structures underwent an energy minimization process using GROMACS. Energy minimization is a crucial step in molecular dynamics simulations to remove any steric clashes or unfavourable interactions that may have arisen during model building or phosphorylation. The steepest descent minimization method was used for this purpose. The steepest descent algorithm iteratively adjusts the atomic positions to reduce the overall potential energy of the system. This method is efficient for removing high-energy overlaps and ensuring that the system reaches a local minimum on the potential energy surface. Both the unphosphorylated wild-type structures and the phosphorylated threonine structures underwent this energy minimization process under identical conditions to ensure consistency.

#### Classification of Localized Regions

Residues of the membrane transporter NRT1.1 were categorized into three groups based on their proximity to the phosphorylated threonine residue: “Phosphosite,” “Near Phosphosite,” and “Away from Phosphosite.” This classification aimed to delineate localized regions within the protein structure to better understand the effects induced by phosphorylation. The “Phosphosite” category included residues within a 4.5 Å radius of the phosphorylated residue, as determined by an interatomic distance cutoff. Residues falling within a range of 4.5 Å to 10 Å from the phosphorylation site were classified as “Near Phosphosite.” Finally, residues located more than 10 Å away from the phosphorylated threonine were designated as “Away from Phosphosite.” This categorization enabled a focused analysis of how phosphorylation influences specific regions of the protein, facilitating a more detailed understanding of the structural and dynamic shifts induced by this post-translational modification.

#### Comparative Conformational Deviation

TM-align method was used to align the structures of phosphorylated and unphosphorylated forms of the membrane transporter [43]. TM-align is a widely recognized structural alignment algorithm in bioinformatics, designed to compare and align protein structures by incorporating both sequential and spatial information. Following the structural alignment, the backbone root-mean-square deviation (RMSD) was computed. RMSD serves as a quantitative measure of structural dissimilarity, representing the average distance between corresponding atoms in two aligned structures. A higher RMSD value indicates greater structural deviation. Mathematically, the RMSD between two sets of coordinates (X1, Y1, Z1) and (X2, Y2, Z2) for a set of atoms is calculated as follows:

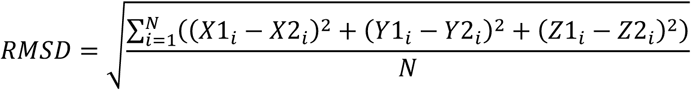

Here, N represents the number of atoms considered in the calculation. To assess local structural changes, individual alpha carbon (Cα) deviations between equivalent residues in the phosphorylated and unphosphorylated forms were also computed.

#### Comparative Dynamics Assessment

Understanding long-timescale protein motions is essential for deciphering their functional significance, with Normal Mode Analysis (NMA) being a favoured method for such studies [20, 21, 44–46]. NMA utilizes Cartesian coordinates from the protein structure and a force field that defines inter-atomic interactions. This method involves creating a ‘Hessian’ matrix from the second derivative of potential energy, which is then diagonalized to obtain eigenvalues and eigenvectors. Global protein motions, particularly low-frequency collective movements, have been shown in previous studies to play a significant role in biological functions. Due to the computational intensity of all-atom NMA, a coarse-grained approach was adopted using the Cα-level Anisotropic Network Model (ANM)-based NMA for this study [47]. The ProDy package was employed for these calculations [48]. In the coarse-grained model, Cα atoms are represented as masses connected by elastic springs with identical spring constants if their inter-Cα distance is less than 15 Å. Contributions from the first 5 N-terminal and last 5 C-terminal residues were excluded from the analysis. Normal modes were computed for NRT1.1 protein pairs in both phosphorylated and unphosphorylated forms. Mean squared fluctuations from the first 10 consecutive non-zero normal modes were scaled using z-score normalization. To quantify the dynamic alterations, the Root Mean Squared Difference of Fluctuations (RMSDf) was computed. RMSDf, analogous to RMSD, measures the difference between the normalized fluctuations of the phosphorylated and unphosphorylated forms. The Overlap, representing the inner product of eigenvectors from the 10 lowest frequency modes, was calculated using the ProDy package, to evaluate the accessibility of intrinsic motions in the phosphorylated form compared to the unphosphorylated form. This metric provided insight into the extent of shared conformational space between the two forms, offering a deeper understanding of the protein’s dynamic behaviour.

#### Modelling Protein-Protein Docked Complexes

In the RCSB Protein Data Bank, no structures are available for the protein-protein complexes of CIPK8 and CIPK23 with CBL1 (i.e., CIPK8-CBL1 complex and CIPK23-CBL1 complex). Therefore, these complexes were modelled in silico using the AlphaFold Server [49]. The FASTA sequences for CIPK8, CIPK23, and CBL1 from Arabidopsis thaliana were obtained from UniProt [50]and submitted to the AlphaFold3 webserver. The top-ranked model, with the highest iPTM and PTM scores, was selected for further analysis.

#### Assessing Interaction Strength and Interfacial Residues

The strength of the protein-protein interactions and the identification of interfacial residues for the CIPK8-CBL1 and CIPK23-CBL1 complexes were estimated using PPCheck [51]. Interaction measurements were based on standard energy calculations involving non-bonded interactions such as van der Waals forces, electrostatic interactions, and hydrogen bonds. Hydrogen bond energies were calculated using Kabsch & Sander’s equation, while electrostatic interaction energy was determined using Coulomb’s equation with a distance-dependent dielectric constant. van der Waals energy calculations included hydrophobic interactions and contributions from van der Waals pairs. Total stabilizing energy was derived from the sum of these non-bonded interactions. Interface residues were identified as those contributing to any non-bonded interactions, with specific criteria for hydrophobic interactions (CB-CB atom distance less than 7Å) and salt bridges (side-chain nitrogen and oxygen atoms of oppositely charged residues within 4Å).

#### Solvent Accessibility and Solvation Energy

The solvent accessible surface area (SASA) and solvation energy of NRT1.1 before and after Threonine 101 phosphorylation were calculated using GetArea [52]. This calculation provides insights into how phosphorylation at Threonine 101 impacts the exposure of residues to the solvent and the overall solvation energy, which is crucial for understanding conformational changes and functional implications.

#### Visualization

Protein structure visualization was performed using UCSF Chimera [53]. To explore normal modes, “.nmd” extension files generated by ProDy were imported into the NMWiz (Normal Mode Wizard) plugin of Visual Molecular Dynamics (VMD) [54]. For comparative assessments of normal modes, videos showcasing backbone fluctuations and capturing the dynamic behaviour of the proteins were produced using the MovieMaker plugin in VMD. These videos were further refined with VideoMach to enhance their presentation quality.

### Systems Analysis

#### Modeling Steady-State Flux of the NRT1.1 Transporter

The NRT1.1 protein transporter facilitates the movement of nitrate ions across the lipid bilayer of the cell membrane by alternately exposing its nitrate binding site to either the extracellular or intracellular environment. This alternating exposure enables nitrate to be transported from one side of the membrane to the other. This model is inspired by previously built models for glucose transporters, where similar transport mechanisms have been well-studied [27, 28].

We can model the NRT1.1 transport process by considering two conformational states: Ci and Ce, where Ci refers to the transporter with the nitrate binding site exposed to the intracellular side, and Ce refers to the transporter with the nitrate binding site exposed to the extracellular side. Since NRT1.1 is a proton-coupled symporter, the nitrate and the proton substrates bind to these transporter states. We denote by Si the intracellular substrate concentration binding to Ci to form the complex Pi, and Se the extracellular substrate concentration binding to Ce to form the complex Pe.

The transport cycle involves transitions between four states: Ce, Pe, Pi, and Ci. These conformational changes are key to the overall transport mechanism. The rate of change of each state can be described by differential equations.

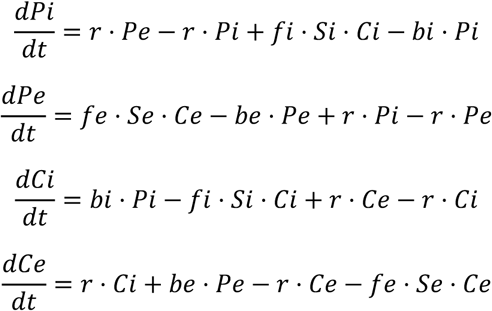

At steady state, the transporter’s flux J is

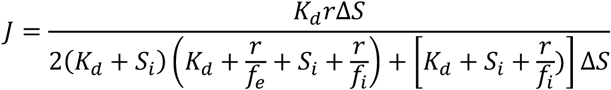

which can further be written as a Michaelis-Menten function of the difference between the extra- and intracellular concentrations of substrate.

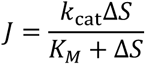

with

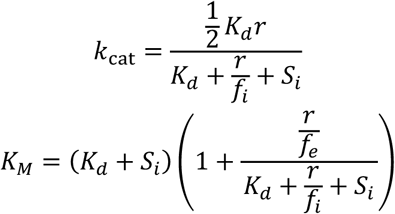

where,

fe: Association rate of extracellular substrate to the transporter

be: Dissociation rate of extracellular substrate to the transporter

fi: Association rate of intracellular substrate to the transporter

bi: Dissociation rate of intracellular substrate to the transporter

r: Transition rate of the transporter across the membrane

A significant finding, based on observations from Parker et al. [13], is that the dissociation constant (*K*_*d*_) for nitrate binding; defined as

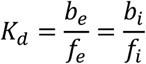

remains largely unchanged before and after phosphorylation. This indicates that the binding affinity for nitrate is unaffected by the phosphorylation state. However, the key parameter r, which governs the transition rates of the transporter across the membrane, reflects changes in the protein’s flexibility. Our results suggest that phosphorylation at Threonine 101 alters this flexibility, making it reasonable to conclude that r likely shifts after phosphorylation. This change in r directly affects the *K*_*M*_ value, which in turn modulates the overall flux (J) of the transporter. In the phosphorylated state, the transition rate is r^phos^, and in the unphosphorylated state, it is r^unphos.^ The steady-state flux (J) will differ between these two states due to this variation in the transition rates.

#### Simulation of Steady-State Flux of the NRT1.1 Transporter

We simulated the transport cycle of the NRT1.1 transporter system with the following assumptions. We first assumed that the ratios be/fe and bi/fi are equal as previously described and were set to 1 corresponding the Kd of the transporter system which was experimentally determined to be 1mM. This value was kept constant for both the unphosphorylated and phosphorylated forms of the protein. The internal concentration of nitrate was assumed to be 1e-5 times the external nitrate concentration that was varied over a range of 0 to 10mM. Furthermore, fi was assumed to be 1e3 times the value of fe. This assumption was based on previously estimated parameters in the context of glucose transporters. The simulated output flux J at steady state of the transporter was calculated as bi*Pi - fi*Si*Ci. Kcat and Km for each parameter set were obtained by fitting the simulated levels of J to a Michaelis-Menten function for different values of Se and Si using the curvefit function in python. The initial condition for Pe was set to 1 while all others were set to 0. This implies that the total of all conformations corresponds to 1 which upon simulation to steady state would result in the probability of observing any of the conformations. The time for each simulation was set till 2000 with computations being done at an interval of 0.2-time units within which all the parameter sets reached steady states. Simulations were performed over ranges of values of r, fi and the ratio between the r of the phosphorylated and unphosphorylated forms using the odeint function in python. For the simulation of the biphasic response of NRT1.1 we assumed that the r parameter changed as a function of the external nitrate concentration as a Hills function with threshold being set at 1.5 and with a hill’s coefficient 10. These parameters are qualitatively assumed currently based on previous data on experimental biphasic response curves. This assumption was replaced by systems level network changes in the final set of simulations of the NRT1.1 transport dynamics along with simulations of the gene regulatory network.

#### Simulation of the NRT1.1 transport dynamics coupled with regulatory network

Simulations of the gene regulation of NRT1.1 and its phosphorylation form were performed by defining a system of coupled differential equations. The key features of the gene regulatory network include – a) activation of the NRT1.1 signaling pathway by external nitrate; b) activation of CIPK8 through due to activation of NRT1.1 via NLP7; c) sequestration of CBL from CIPK23 due to higher abundance of CIPK8; d) reduced CIPK23-CBL based phosphorylation of NRT1.1 leading to increase in unphosphorylated form of NRT1.1. The following equations were used to simulate the above set of regulatory interactions.

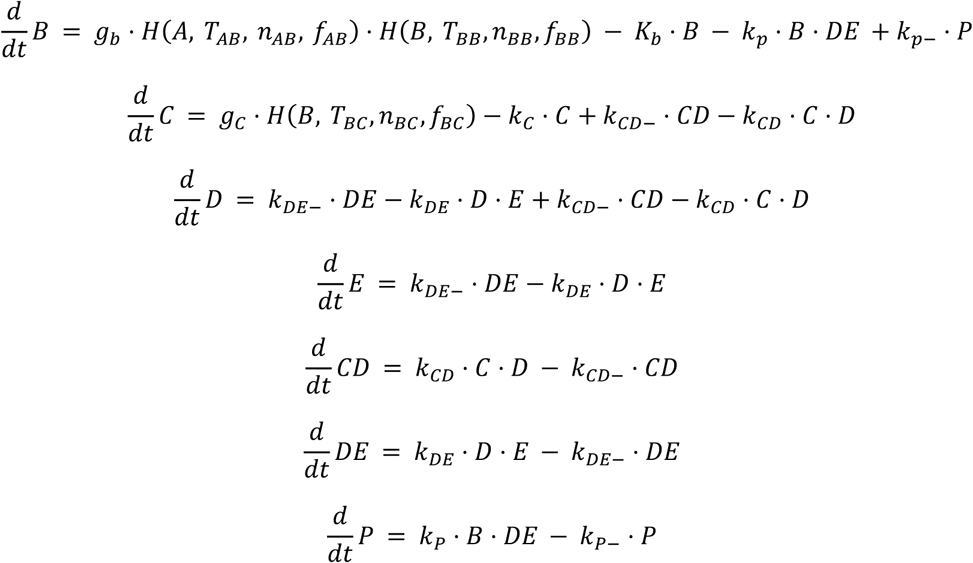

In the above set of equations, B represents the unphosphorylated form of NRT1.1, C is CIPK8, D is the amount of free CBL, E is CIPK23, CD is the amount of CIPK8-CBL complex, DE is the amount of CIPK23-CBL complex, and P is the phosphorylated form of NRT1.1. H represents the shifted Hills function given by the following expression

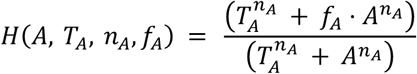

This function approaches f_A_ as A tends to infinity and is 1 if A = 0. The hills coefficient controls the steepness of the curve.

The description of the parameters and the corresponding parameter values are provided in the **Supplementary Table**. Most of the parameter values are assumed parameter values are estimation of each parameter does not exist in the literature. As a result, our simulation results show a possible proof of concept mapping to the known biology rather than a quantitative simulation of the real simulation. The above set of differential equations were simulated till they reached steady states and the steady state values were reported in the graphs. The initial conditions used to simulate the above set of differential equations are also listed in the **Supplementary Table**. All the simulations were performed in python using *odeint*.

## Author Contributions

SSu: conceptualized and conducted research, including structural simulations, kinetic modeling, system-level regulatory network design and simulation, and prepared the first draft of the manuscript. SSa: contributed to kinetic and dynamic simulations of system-level regulatory network. MR: Conceptualized and supervised research and edited the manuscript. MKJ: Analysed data and edited the manuscript. MR and MKJ acquired the funding.

## Acknowledgement

This work is supported by the Department of Science and Technology, Government of India [Grant No. DST/INSPIRE/04/2020/001492] and the Science and Engineering Research Board, Government of India [Grant No. CRG/2023/006432] to Mubasher Rashid. M.K.J. was supported by Ramanujan Fellowship (SB/S2/RJN-049/2018), Science and Engineering Board, Government of India. M.K.J. was also supported by Param Hansa Philanthropies.

## Code Availability

The supplementary movie file and python scripts are available here: https://github.com/seemadri/Kinetics_NitrateUptake_NRT1.1

## Supplementary Figures

**Supplementary Figure 1:**
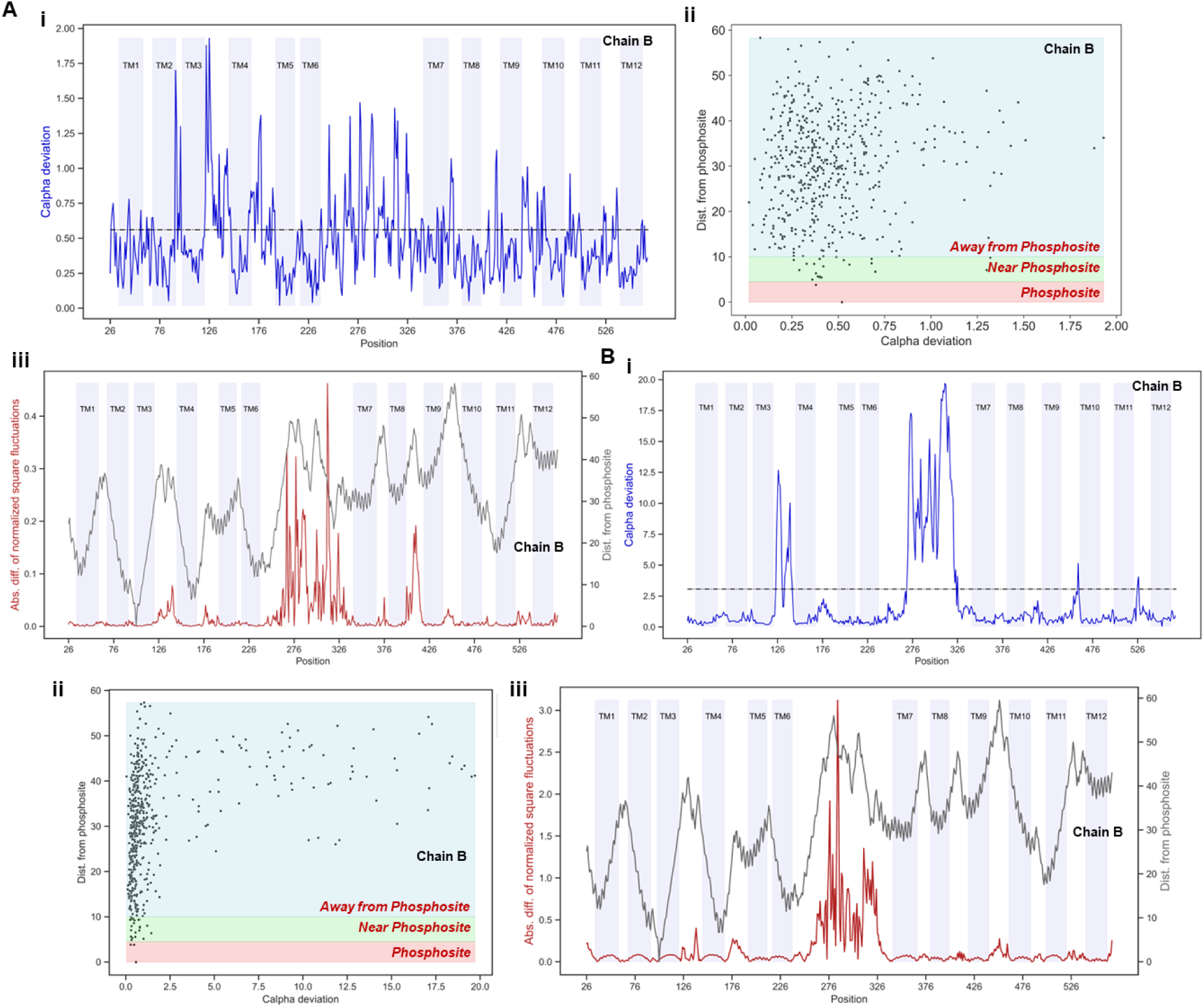
Cα deviations and absolute differences in normalized square fluctuations in the second chain of the NRT1.1 homodimer. **(A) (i)** Cα deviations between the Threonine 101 phosphorylated and apo forms of NRT1.1 for all aligned residues (Chain B), indicated by a blue line. The overall root mean square deviation (RMSD) between these two forms is denoted by a black line. Regions corresponding to transmembrane helices (TM1 to TM12) are coloured lavender. **(ii)** Scatterplot showing Cα deviations versus the distance of corresponding residues from the phosphosite. Regions labeled as “Phosphosite,” “Near Phosphosite,” and “Away from Phosphosite” are highlighted in red, green, and blue shades, respectively. **(iii)** Absolute difference between normalized square fluctuations of the Threonine 101 phosphorylated and apo forms of NRT1.1 for all aligned residues (Chain B), depicted by a maroon line. The grey line represents the distance of each residue from the phosphorylation site. **(B) (i)** Cα deviations between the phosphorylated nitrate-bound and apo forms of NRT1.1 for all aligned residues (Chain B), represented by a blue line. The overall RMSD between these two forms is shown by a black line. **(ii)** Scatterplot illustrating the relationship between Cα deviations and the distance of corresponding residues from the phosphosite. Regions labeled as “Phosphosite,” “Near Phosphosite,” and “Away from Phosphosite” are marked in red, green, and blue shades, respectively. **(iii)** Absolute difference between normalized square fluctuations of the nitrate-bound Threonine 101 phosphorylated and apo forms of NRT1.1 for all aligned residues (Chain B), depicted by a maroon line. The grey line represents the distance of each residue from the phosphorylation site.

**Supplementary Figure 2:**
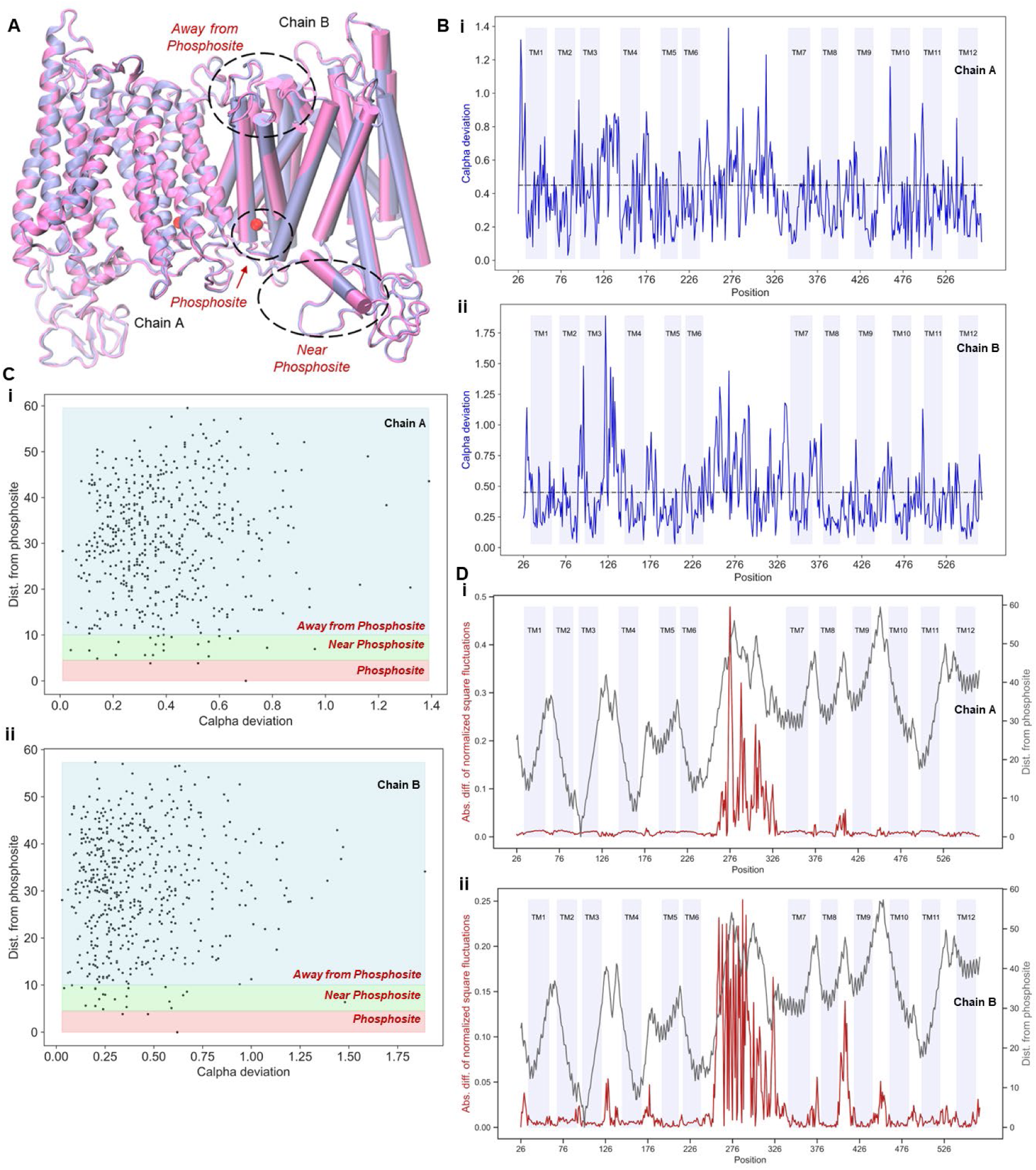
Comparative Structural Dynamics Analysis of NRT1.1 bound to nitrate (PDB ID: 5A2O) and its in-silico modelled Threonine 101 phosphorylated forms. **(A)** Structural superimposition of the nitrate-bound form and its Threonine 101 phosphorylated counterpart of NRT1.1, with the phosphorylated form coloured pink. Black circles highlight regions showing structural disparities, including the phosphosite, areas near the phosphosite, and regions further away. The phosphorylated residue is represented by a red sphere. **(B)** Cα deviations between the phosphorylated nitrate-bound and unphosphorylated nitrate-bound forms of NRT1.1 for all aligned residues, depicted for **(i)** Chain A and **(ii)** Chain B, with deviations shown by a blue line. The overall RMSD between these two forms is represented by a black line. **(C)** Scatterplot illustrating the relationship between Cα deviations and the distance of corresponding residues from the phosphosite, shown for **(i)** Chain A and **(ii)** Chain B. Regions labelled “Phosphosite,” “Near Phosphosite,” and “Away from Phosphosite” are shaded in red, green, and blue, respectively. **(D)** Absolute difference in normalized square fluctuations between the nitrate-bound Threonine 101 phosphorylated and unphosphorylated forms of NRT1.1 for all aligned residues, shown for **(i)** Chain A and **(ii)** Chain B, depicted by a maroon line. The grey line indicates the distance of each residue from the phosphorylation site.

**Supplementary Figure 3:**
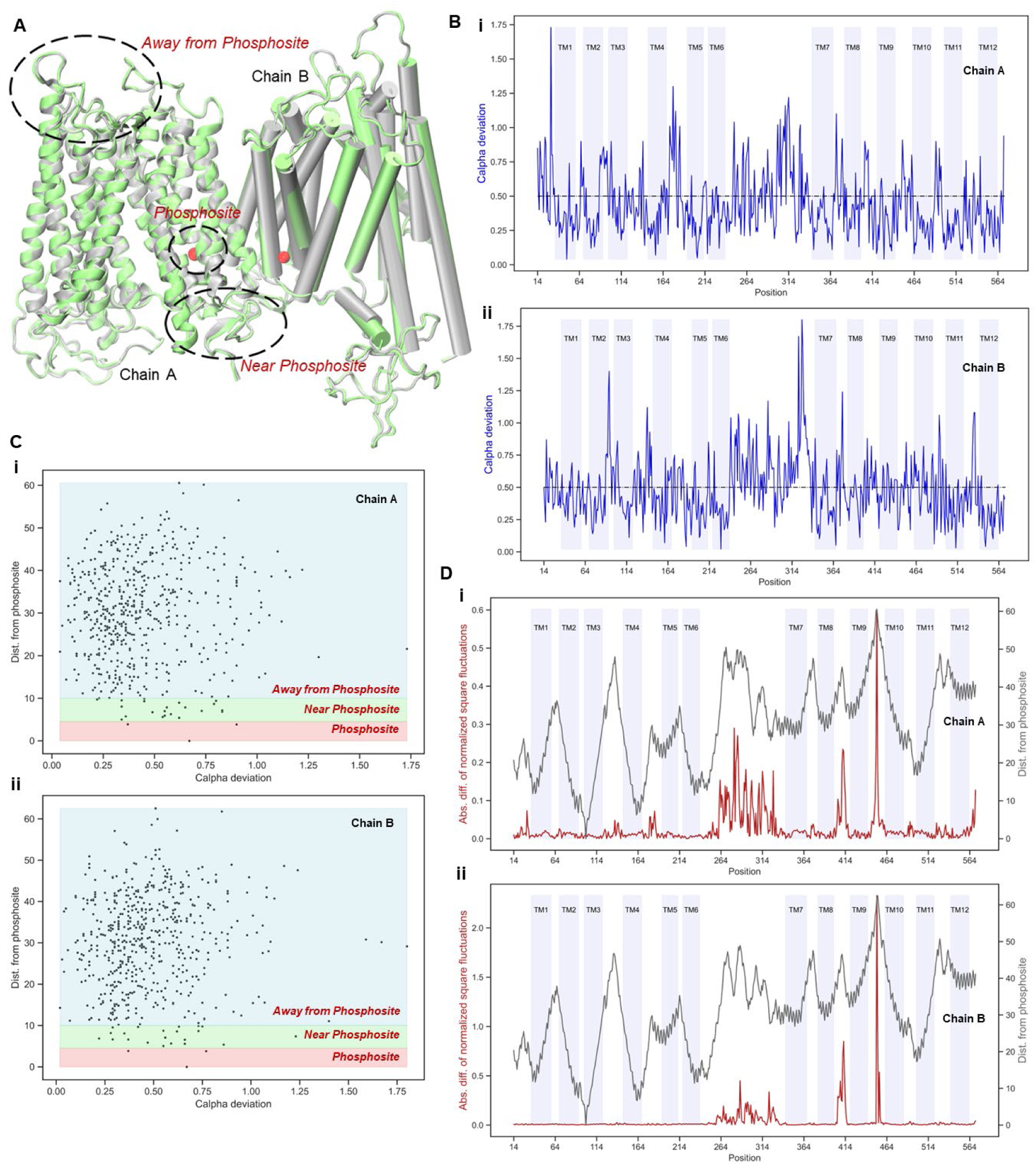
Comparative Structural Dynamics Analysis of NRT1.1 bound to nitrate (PDB ID: 4OH3) and its in-silico modelled Threonine 101 phosphorylated forms. **(A)** Structural superimposition of the nitrate-bound form and its Threonine 101 phosphorylated counterpart of NRT1.1, with the phosphorylated form coloured green. Black circles highlight regions exhibiting structural disparities, including the phosphosite, nearby areas, and regions farther away. The phosphorylated residue is depicted as a red sphere. **(B)** Cα deviations between the phosphorylated nitrate-bound and unphosphorylated nitrate-bound forms of NRT1.1 for all aligned residues, shown for **(i)** Chain A and **(ii)** Chain B, with deviations represented by a blue line. The overall RMSD between these forms is indicated by a black line. **(C)** Scatterplot illustrating the relationship between Cα deviations and the distance of corresponding residues from the phosphosite, shown for **(i)** Chain A and **(ii)** Chain B. Regions labelled “Phosphosite,” “Near Phosphosite,” and “Away from Phosphosite” are shaded in red, green, and blue, respectively. **(D)** Absolute difference in normalized square fluctuations between the nitrate-bound Threonine 101 phosphorylated and unphosphorylated forms of NRT1.1 for all aligned residues, shown for **(i)** Chain A and **(ii)** Chain B, depicted by a maroon line. The grey line represents the distance of each residue from the phosphorylation site.

